# SICKO: Systematic Imaging of *Caenorhabditis* Killing Organisms

**DOI:** 10.1101/2023.02.17.529009

**Authors:** Luis S. Espejo, Samuel Freitas, Vanessa Hofschneider, Leah Chang, Angelo Antenor, Jonah Balsa, Anne Haskins, Destiny DeNicola, Hope Dang, Sage Hamming, Delaney Kelser, George L. Sutphin

## Abstract

*Caenorhabditis elegans* are an important model system for research on host-microbe interaction. Their rapid life cycle, short lifespan, and transparent body structure allow simple quantification of microbial load and the influence of microbial exposure on host survival. *C. elegans* host-microbe interaction studies typically examine group survival and infection severity at fixed timepoints. Here we present an imaging pipeline, Systematic Imaging of *Caenorhabditis* Killing Organisms (SICKO), that allows longitudinal characterization of microbes colonizing isolated *C. elegans*, enabling dynamic tracking of tissue colonization and host survival in the same animals. Using SICKO, we show that *Escherichia coli* or *Pseudomonas aeruginosa* gut colonization dramatically shortens *C. elegans* lifespan and that immunodeficient animals lacking *pmk-1* are more susceptible to colonization but display similar colony growth relative to wild type. SICKO opens new avenues for detailed research into bacterial pathogenesis, the benefits of probiotics, and the role of the microbiome in host health.

## Introduction

*Caenorhabditis elegans* are an important model system for host-microbe research due to the ability to rapidly quantify the influence of microbial exposure on whole-organism survival and estimate microbial load. Immune research using mammalian model systems often relies on *in vitro* studies in which cell types are isolated to simulate the *in vivo* environment using cell culture, or in whole animal studies where the initiation and progression of microbial interaction is difficult to quantify and time consuming (Bar-Ephraim et al., 2020; Wagar et al., 2018). Invertebrate organisms offer the ability to study the immune system in whole organisms with a short lifespan at relatively high throughput. The roundworm *Caenorhabditis elegans* lacks adaptive and humoral immune components present in mammals but recapitulates many genetic and molecular elements of immunity shared across the animal kingdom (Bar-Ephraim et al., 2020; Fabian et al., 2021). Because *C. elegans* have a transparent body cavity, microbes may be easily observed when they colonize the tissues of individual animals (Elkabti et al., 2018; Marsh and May, 2012; Rutter et al., 2019; Tan et al., 1999).

Using current standard approaches, *C. elegans* are typically anesthetized to visualize tissue colonization by fluorescently labeled bacteria, which results in animal death and limits observations to a cross-section of the population at a specified timepoint. This precludes observation of dynamic changes in colonization of individual animals, such as timing of infection initiation, colony growth rate, and the potential reversal of colony growth—in individual animals (Boulin et al., 2008; Massie et al., 2003; Shaham, 2006). This also prevents measurement of survival or other metrics on of long-term health in the individual animals for which colony size and location was measured. For this reason, the relationship between infection dynamics and physiological outcomes, including long term health and survival, is not well understood.

Here we present a new system called Systematic Imaging of *Caenorhabditis* Killing Organisms (SICKO) capable of characterizing longitudinal interactions between host and microbes in individual *C. elegans*. We recently reported a new method for long-term cultivation of *C. elegans* that is conducive to longitudinal fluorescent imaging of individual *C. elegans* isolated on solid nematode growth media (NGM) (Espejo et al., 2022). SICKO leverages this culture system to track microbial colonization in the tissues of individual free-crawling *C. elegans*, enabling researchers to capture dynamic changes in gut colonization and assess the subsequent impact on health and survival in the same animals. Using this system, we demonstrate that gut colonization by *Escherichia coli* strain OP50, commonly used as a laboratory food source, dramatically shortens *C. elegans* lifespan. Validation experiments demonstrate that immunodeficient animals lacking the *pmk-1* gene do not display altered progression of bacterial colony growth, but rather suffer an increased rate of gut colony initiation. Finally, we show that the gram-negative pathogen *Pseudomonas aeruginosa* strain PA14 displays increased infectability, toxicity, pathogenicity, and infection progression relative to *E. coli* OP50. SICKO provides a powerful tool into understanding the mechanisms of host-microbe interaction, opening new avenues for detailed research into therapies that combat pathogen induced illness, the benefits imparted by probiotic bacteria, and the role of the microbiome in host health.

## Results

### SICKO tracks fluorescently labeled bacteria colonies in individual C. elegans

Each SICKO experiment begins by age-synchronizing a population of *C. elegans* using standard techniques and initially maintaining animals in group culture on petri plates under standard conditions (NGM seeded with *E. coli* food; **Fig. 1a[i]**) (Porta-de-la-Riva et al., 2012). At a user-specified age, animals are exposed to bacteria modified to constitutively express a fluorescent protein (**Fig. 1a[ii]**). The length of exposure should be sufficient to allow gut colonization in a subset of the population and must be optimized for each bacterial strain of interest (e.g., 7 days starting at the L4 larval stage for *E. coli* strain OP50; 2 days starting at day 5 of adulthood for *P. aeruginosa* strain PA14). Following the exposure period, we move worms to petri plates containing NGM seeded with non-fluorescent bacteria for at least 16 hours to allow non-adherent bacteria to be passed from the gut and external bacteria to detach from the cuticle (**Fig. 1a[iii]**). Step iii ensures that the only fluorescently labeled bacteria remaining is that which has colonized the animal, as signal from non-colonizing bacteria on the animal or on the plate will generate signal that will confound the quantification of fluorescent colonies adherent to the internal tissues of the worm (**Fig 1b**). Worms are then transferred to individual wells in a culture environment designed to isolate animals on NGM pads seeded with non-fluorescent bacteria (**Fig. 1a[iv], c**), the preparation of which we recently published (Espejo et al., 2022). Finally, fluorescent images of individual free-crawling worms are captured daily, and worms are manually scored each day for survival and fleeing (**Fig. 1a[v]**). Once image collection is complete for an entire experiment, images are analyzed to quantify the area and intensity of fluorescent bacteria in the gut at each time point (**Fig. 1a[vi]**).

**Figure 1.**
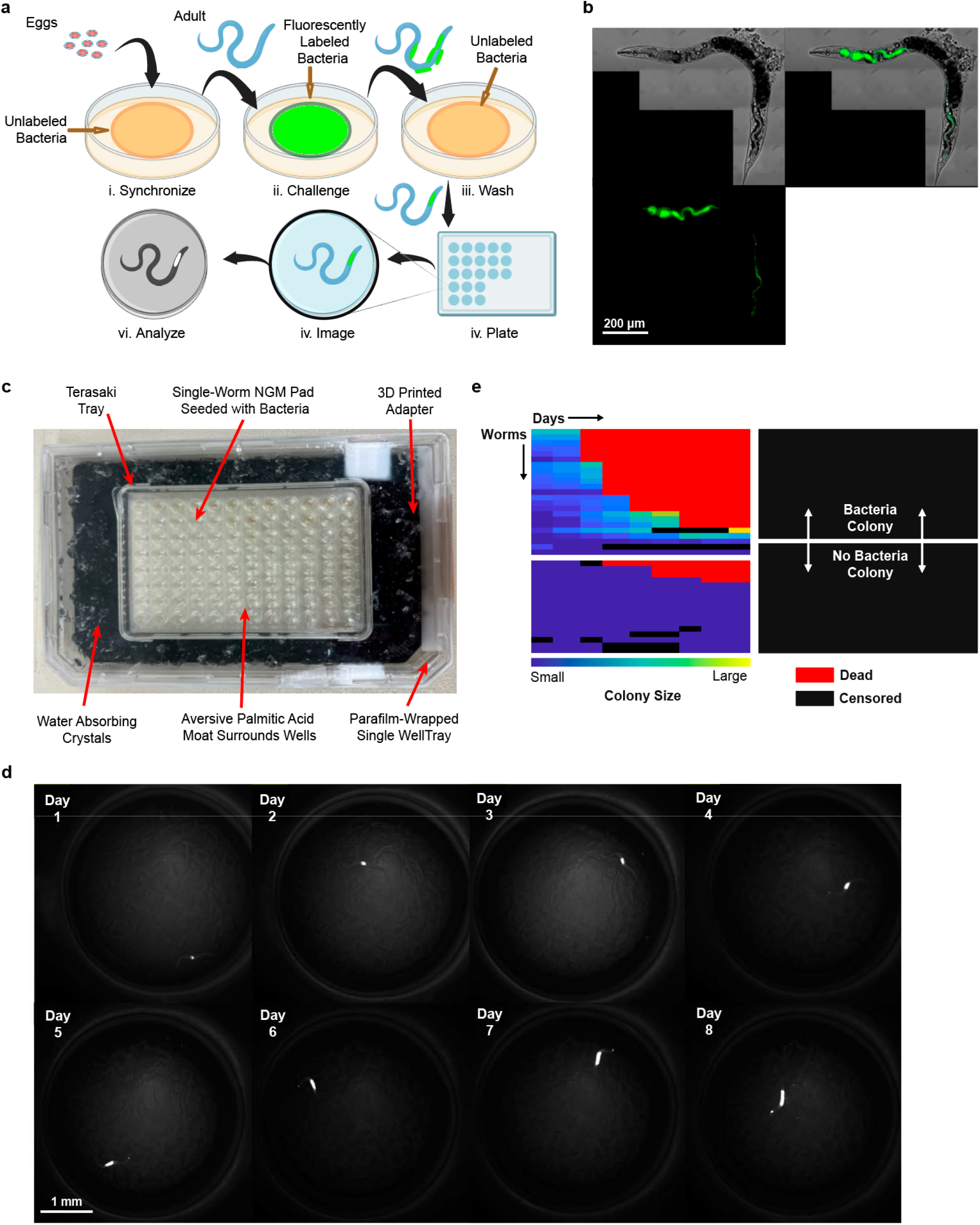
Overview of SICKO workflow and data output. (**a**) Flow chart of SICKO workflow. [i] *C. elegans* are age-synchronized as eggs and allowed to grow to adulthood. [ii] Adult worms are challenged with fluorescently labeled bacteria or other microbes. [iii] Following challenge, worms are allowed to crawl on media seeded with non-fluorescent bacteria for 16 hours to wash bacteria from intestine and external surfaces that are not part of an adherent colony. [iv] Individual animals are plated on wells within a single-worm culture environment and [v] imaged daily. [vi] Images are analyzed and compiled. (**b**) Representative confocal image of an individual *C. elegans* harboring an intestinal *E. coli* labeled with GFP following challenge and wash. (**c**) Image of a single-worm culture environment. Worms are housed on individual wells of a Terasaki tray filled with NGM and seeded with bacteria. Wells are surrounded by an aversive barrier of palmitic acid to prevent fleeing. The Terasaki tray is mounted inside a single-well tray on a custom 3D printed adapter. The space around the Terasaki tray is filled with saturated water absorbing crystals, and the single well tray is wrapped in parafilm to prevent the wells from drying out. (**d**) Representative images of a single *C. elegans* harboring a GFP labeled *E. coli* intestinal colony over time. Images were taken using the GFP channel on a fluorescent stereoscope and processed with the SICKO software. GFP-labeled *E. coli* is shown in white. (**f**) Representative heatmap representation of data from a population of *C. elegans* challenged with fluorescently labeled bacteria. Each row is a single animal and each column is a day. Blue and yellow boxes indicate colony size. Red boxes indicate dead worms and black boxes indicate censored data. Rows above the white line represent animals harboring a bacteria colony, while rows below the white line represent animals without a colony.

SICKO results in a rich dataset capturing colony progression throughout life and lifespan for each animal. The single-worm culture environment isolates individual *C. elegans* on small NGM pads surrounded by an aversive barrier of palmitic acid to discourage animals from fleeing their well (**Fig. 1c**). This distinguishes SICKO from methods that use microfluidic systems to monitor bacterial colonization in individual animals (Viri et al., 2021), allowing a more direct comparison to the majority of previous studies that use *C. elegans* to examine host-microbe interactions on solid media. The culture environment is enclosed in a chamber humidified with water absorbing beads, allowing for long-term cultivation of animals with repeated sampling of colony data and measurement of lifespan (**Fig. 1c**). The output of SICKO provides detailed information about when a bacterial colony becomes large enough to be detectable, colony size, and colony growth progression for each individual animal (**Fig. 1d**), in addition to insights into population-level dynamics such as infectivity and the impact of bacterial colonization and growth on host health and survival. The infection life history data for a population can be summarized in a heat map that reflects daily infection area or intensity for each animal, survival, and censoring events (**Fig. 1e**). Because the animals are imaged while crawling freely, the quantitative longitudinal infection data generated by SICKO comes with the current limitation of identifying the precise location of the infection within the worm. Each single-worm culture environment houses 96 animals and can be scaled using additional environments. In addition to allowing observation of the dynamic growth of individual colonies and the direct correlation of these dynamics to survival and health in the same animal, isolating individual animals also increases efficiency by allowing sample size to be precisely tuned to the needs of each experiment.

### SICKO captures bacterial pathogen sensitivity of C. elegans lacking pmk-1

To validate SICKO, we first monitored *C. elegans* deficient in the p38 MAPK ortholog *pmk-1*, a central regulator of the *C. elegans* innate immune response to pathogenic bacteria (Engelmann and Pujol, 2010; Pukkila-Worley and Ausubel, 2012; Shivers et al., 2010; Troemel et al., 2006), challenged with *E. coli* strain OP50. *E. coli* OP50 is the strain most commonly used as a *C. elegans* laboratory food source and is thought to be mildly pathogenic. Specifically, worms fed UV-inactivated *E. coli* OP50 are longer-lived (Garigan et al., 2002; Gems and Riddle, 2000; Win et al., 2013), while worms fed live *E. coli* OP50 grown on alternative media that promotes bacterial growth are shorter-lived (Garsin et al., 2001; Mylonakis et al., 2002), than worms fed live *E. coli* OP50 grown on NGM. *pmk-1* mutants have reduced lifespan relative to wild type when cultured on live *E. coli* OP50 (Alper et al., 2010).

We exposed animals to *E. coli* OP50 constitutively expressing green fluorescent protein (GFP) under a trc promoter for 7 days starting at the L4 larval stage and monitored infection. SICKO detected significantly higher rates of colonization in *pmk-1* mutants relative to wild type (**Fig. 2a, S1a**). Colonization significantly correlated with death for both strains (**Fig. 2a**). To compare the SICKO output to cross-sectional data generated in previous studies, we simulated a cross-sectional study design by only analyzing colonization on day 1 following challenge (16 hours after wash; **Fig. 1a[iii]**). The proportion of each population with a detectable colony (**Fig. 2b**), integrated infection intensity within each animal (**Fig. S1b**), and area of infection within each animal (**Fig. 2c**) were not significantly different for *pmk-1* mutants relative to wild type using this cross-sectional approach. The cross-sectional design missed ∼29% of colonies that were present but below the detection threshold one day after challenge, but later grew into detectable colonies (**Fig. 2d**). Using SICKO we were able to capture late-emerging colonies through longitudinal imaging, and *pmk-1* animals ultimately displayed a significantly higher total colonization rate (**Fig. 2d**). The day when a colony first becomes detectable significantly correlated with the day of death following challenge in both the wild-type and *pmk-1* mutants (**Fig. 2e, f**), and time between detection of a colony and the death of an animal was slightly but significantly longer in *pmk-1* relative to wild type animals (**Fig. 2g**). In summary, *pmk-1* mutant animals display a higher rate of colonization than wild type animals, though this is only apparent after allowing colonies initially below the detection threshold to progress for several days. SICKO is capable of capturing this pattern, while a single examination of cross-section colonization rates using standard timing did not.

**Figure 2.**
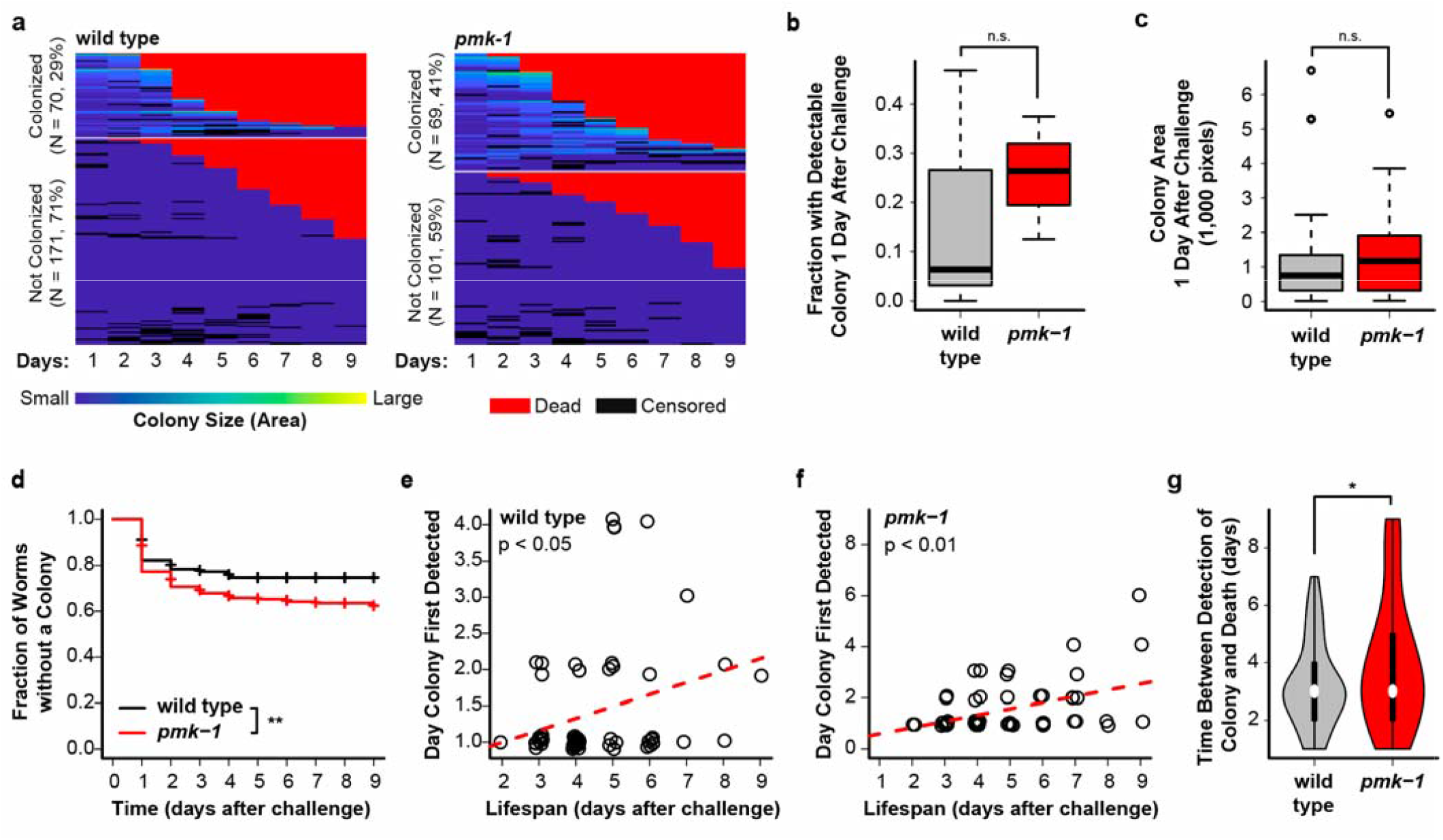
*C. elegans* lacking *pmk-1* are more susceptible to *E. coli* colonization than wild type. (**a**) Heatmap representing colony size (based on fluorescent area) for wild type (left) and *pmk-1* mutant (right) *C. elegans* challenged with GFP-labeled *E. coli*. Death was highly correlated with colonization status in both wild type (p < 0.001, Pearson’s chi-squared test) and *pmk-1* (p < 0.001, Pearson’s chi-squared test) animals, and with genotype (p < 0.01, Pearson’s chi-squared test). (**b**) The proportion of animals harboring a detectable *E. coli* colony (p = 0.67, two-sided Welch’s t test) and (**c**) mean colony area (p = 0.36, two-sided Welch’s t test) were not significantly different between wild type and *pmk-1* knockout *C. elegans* 1 day following challenge. (**d**) Animals lacking *pmk-1* developed significantly more late emerging colonies that were not initially detectable that wild type animals (p < 0.01, log rank test). In *C. elegans* harboring an *E. coli* colony, the day when the colony was first detected was significantly associated with lifespan in both (**e**) wild type (p < 0.05, linear regression) and (**f**) *pmk-1* knockout (p < 0.001, linear regression) animals. Points represent values for individual animals. (**g**) The time between the initial detection of a colony and death was significantly longer for *pmk-1* relative to wild type animals (p < 0.05, Welch’s t test). For box-and-whisker and violin plots, center bar or white point represent median, boxes represent upper and lower quartile, whiskers represent the 5^th^ and 95^th^ percentile, and points indicate outliers. Sample sizes: wild type, N_colonized_ = 70, N_uncolonized_ = 171, N_total_ = 241; *pmk-1*, N_colonized_ = 69, N_uncolonized_ = 101, N_total_ = 170. * p < 0.05, ** p < 0.01, *** p < 0.001, n.s. = not significant for indicated statistical test.

### SICKO captures high pathogenesis of P. aeruginosa

In addition to capturing differences in dynamics of host-microbe interaction between *C. elegans* with different genotypes, SICKO can capture differences in colonization dynamics between different microbes. To validate this capability, we compared *C. elegans* challenged with the *E. coli* OP50 expressing GFP described above to *C. elegans* challenged with *P. aeruginosa* strain PA14, a commonly used gram-negative bacterium that is strongly pathogenic to *C. elegans*, constitutively expressing mScarlet under a trc promoter. As expected, animals challenged with *P. aeruginosa* developed colonies at a higher rate and with increased severity relative to animals challenged with *E. coli* (**Fig. 3a**). Colonization by both bacteria was highly associated with death by 9 days post challenge (**Fig 3a**). Unlike our comparison between wild type and *pmk-1* worms challenged with *E. coli*, the cross-sectional equivalent of observing the animals 16 hours after wash did show a significantly higher number of animals with a colony (**Fig. 3b**) and larger average colony area (**Fig. 3c**) for *P. aeruginosa* relative to *E. coli*. We detected the majority of colonies for both bacteria 16 hours following wash, and SICKO continued to detect new colonies throughout the imaging period in both populations (**Fig 3d**). Not surprisingly, the day when a colony becomes detectable in both *E. coli* and *P. aeruginosa* predicted the day when each animals died (**Fig. 3e, f**). Surprisingly, we observed a longer period between detection of the colony and death for the strongly pathogenic *P. aeruginosa* than the weakly pathogenic *E. coli* (**Fig 3g**). We speculate that this may be a consequence of the ability of *P. aeruginosa* to colonize healthier animals while *E. coli* acts more as an opportunistic pathogen that is only able to colonize less healthy animals.

**Figure 3.**
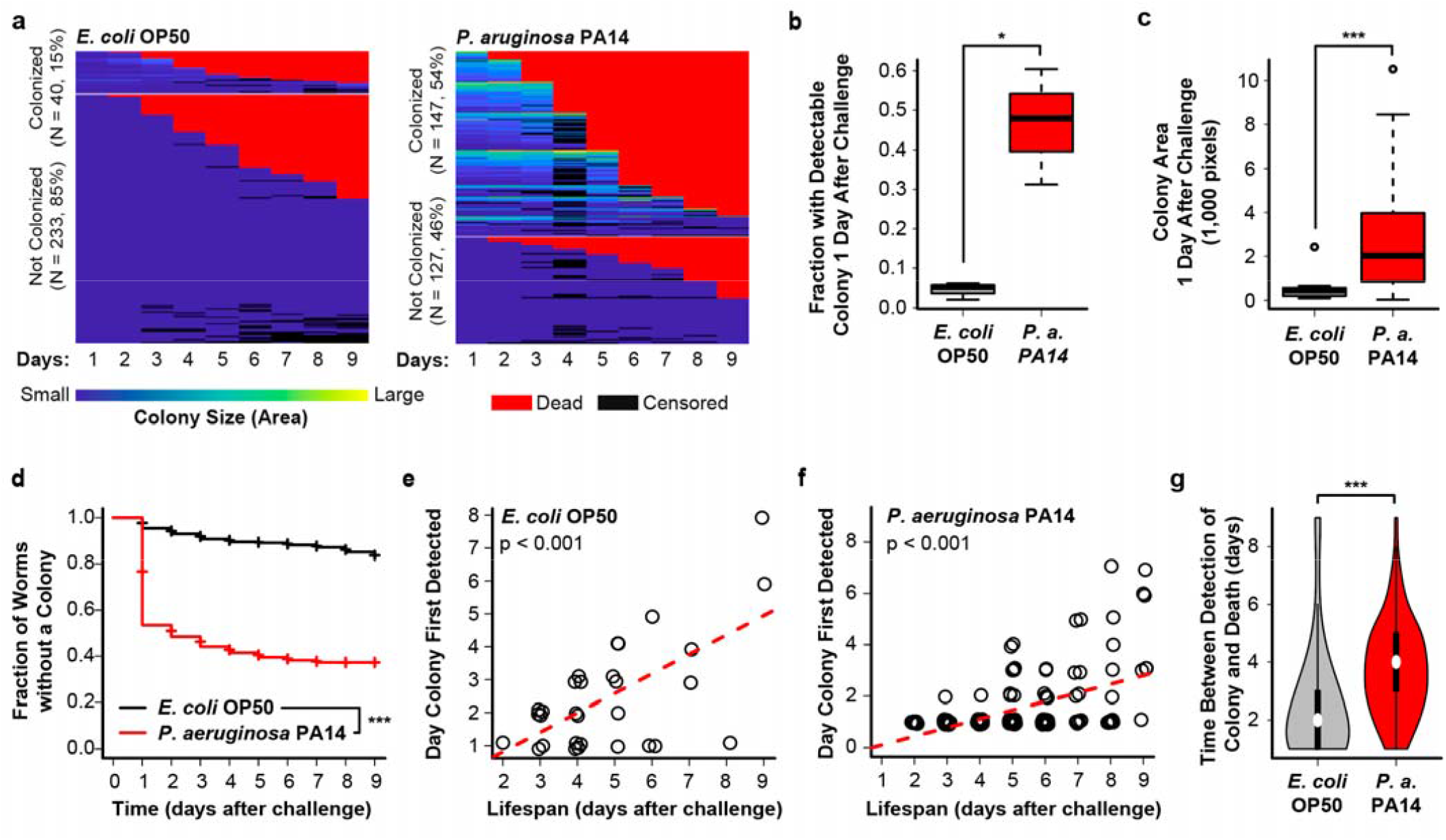
Challenging *C. elegans* with *P. aeruginosa* PA14 results in a greater number of more rapidly progressing colonies relative to *E. coli* OP50. (**a**) Heatmap representing colony size (based on fluorescent area) for wild type *C. elegans* challenges with GFP-labeled *E. coli* OP50 (left) or mScarlet labeled *P. aeruginosa* PA14 (right). Death was highly correlated with colonization status for both *E. coli* (p < 0.001, Pearson’s chi-squared test) and *P. aeruginosa* (p < 0.001, Pearson’s chi-squared test), and with bacterial species (p < 0.001, Pearson’s chi-squared test). (**b**) The proportion of animals harboring a detectable colony (p < 0.05, two-sided Welch’s t test) and (**c**) mean colony area (p = 0.36, two-sided Welch’s t test) were significantly higher for *P. aeruginosa* than *E. coli* 1 day following challenge. (**d**) Animals challenged with *P*. aeruginosa developed substantially more late emerging colonies that were not initially detectable than animals challenged with *E. coli* (p < 0.001, log rank test). In *C. elegans* harboring a colony, the day when the colony was first detected was significantly associated with lifespan for (**e**) *E. coli* (p < 0.001, linear regression) and (**f**) *P. aeruginosa* (p < 0.001, linear regression) challenged animals. Points represent values for individual animals. (**g**) The time between the initial detection of a colony and death was significantly longer for *P. aeruginosa* relative to *E. coli* challenged animals (p < 0.001, two-sided Welch’s t test). For box-and-whisker and violin plots, center bar or white point represent median, boxes represent upper and lower quartile, whiskers represent the 5^th^ and 95^th^ percentile, and points indicate outliers. Sample sizes: *E. coli* OP50, N_colonized_ = 40, N_uncolonized_ = 233, N_total_ = 273; *P. aeruginosa* PA14, N_colonized_ = 147, N_uncolonized_ = 127, N_total_ = 247. * p < 0.05, ** p < 0.01, *** p < 0.001, n.s. = not significant for indicated statistical test.

### SICKO links bacterial colony characteristics to host animal health

*E. coli* OP50 is one of the most utilized laboratory *C. elegans* food sources and its pathogenicity is thought to minimally impact the survival or health of animals with a normal immune system. One application of the SICKO system is to differentiate colonized vs. non-colonized animals within each population. Surprisingly, the survival of both *pmk-1* mutants and wild-type animals was dramatically lower in animals with active OP50 colonies relative to animals without detectable colonies (**Fig. 4a, b**). Additionally, *pmk-1* mutants are thought to be deficient in several aspects of health even in the absence of a pathogen, contributing to the shortened lifespan (Alper et al., 2010). We observed that wild type and *pmk-1* animals with and without detectable OP50 colonization have similar survival (**Fig. 4c**). The SICKO data paints a more complete picture of the health deficits faced by *pmk-1* mutants. It appears that animals lacking *pmk-1* have similar health characteristics to wild type in both colonized and uncolonized subpopulations, but are more likely to be colonized, resulting in a lower survival on average in the combined population.

**Figure 4.**
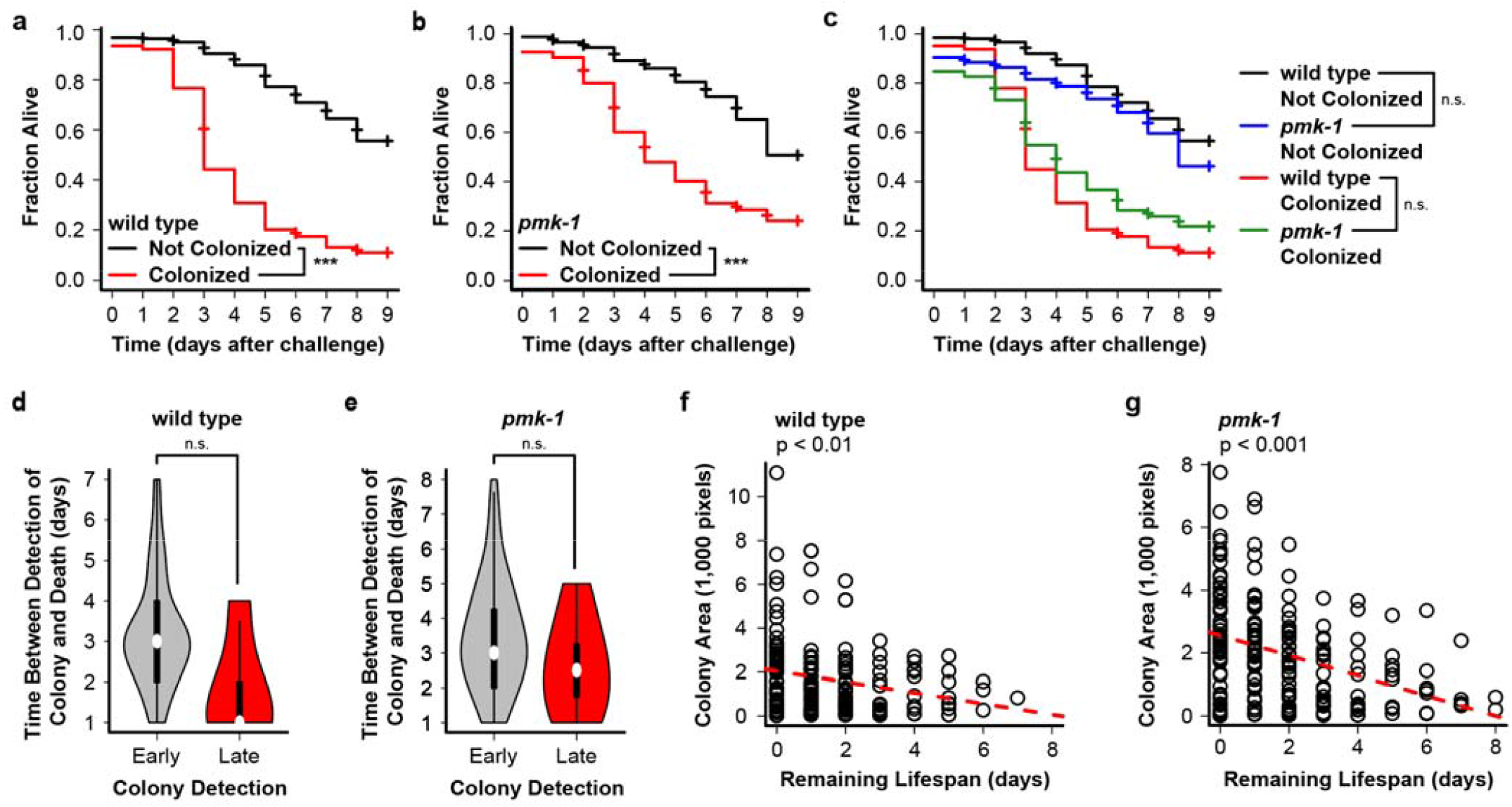
*C. elegans* lacking *pmk-1* have similar lifespan to wild type animals in both colonized and uncolonized subgroups. *C. elegans* harboring a GFP-labeled *E. coli* OP50 colony are substantially shorter lived than animals without colonies in both (**a**) wild type (p < 0.001, log rank test) and (**b**) *pmk-1* knockout populations (p < 0.001, log rank test). (**c**) Lifespan of wild type and *pmk-1* knockout animals is not significantly different within the colonized (p = 0.15, log rank test) and uncolonized (p = 0.055, log rank test) subpopulations. The time between initial detection of a colony and death of the animal trended lower but was not significantly different between animals with colonies that were detected late (day 3 or later post challenge) vs. early (day 1 or 2 post-challenge) for both (**d**) wild type (p = 0.070, two-side Welch’s t test) and (**e**) *pmk-1* (p = 0.13, two-side Welch’s t test) animals. For *C. elegans* harboring an *E. coli* colony, the area of the colony on a given day was significantly and negatively associated with remaining lifespan in both (**f**) wild type (p < 0.01, linear regression) and (**g**) *pmk-1* (p < 0.001, linear regression) animals. Each point represents one animal on one day. For violin plots, center bar or white point represent median, boxes represent upper and lower quartile, whiskers represent the 5^th^ and 95^th^ percentile, and points indicate outliers. Sample sizes: wild type, N_colonized_ = 70, N_uncolonized_ = 171, N_total_ = 241; *pmk-1*, N_colonized_ = 69, N_uncolonized_ = 101, N_total_ = 170. * p < 0.05, ** p < 0.01, *** p < 0.001, n.s. = not significant for indicated statistical test.

These data highlight the ability of SICKO to characterize biological phenotypes in relationship to host-microbe interaction. In both wild type and *pmk-1* worms, the time between a colony being detected and death trended lower for animals with late-detected colonies vs. early-detected colonies, but did not reach significance (**Fig. 4d, e**). Our earlier observation indicates that death typically occurs an average of 3 days after the day when a colony first becomes detectable (**Fig. 2g**). Finally, SICKO allows size of a colony on each day to be isolated and independently compared to remaining survival. We observe a correlation between colony area (**Fig. 4f, g**) or intensity (**Fig. S2a, b**) and remaining lifespan in both wild type and *pmk-1* animals. This provides a framework for predicting the remaining lifespan of an animal at any point in life based on colony size.

When we made these same comparisons between different bacteria, animals with intestinal colonies were again shorter lived than animals without colonies for both *E. coli* (**Fig. 5a**) and *P. aeruginosa* (**Fig. 5b**). Animals challenged with *P. aeruginosa* were shorter-lived than animals challenged with *E. coli* in both the colonized and uncolonized sub-groups (**Fig. 5c**), likely reflecting the fact that part *P. aeruginosa* exert toxicity to their host animals through both colonization and production of excreted toxins that can affect host independent of colonization (Liu, 1974). We also observed that the time between first detection of a colony and animal death was longer for *P. aeruginosa* than *E. coli* for both early- and late-detected colonies (**Fig. 5e, f**). Speculatively, this may again be a consequence of *E. coli* being more of an opportunistic pathogen, selectively colonizing animals with a lower health status, while *P. aeruginosa* initially colonizes healthier animals resulting in a longer period of active colonization before the animals dies. Finally, unlike our first set of experiments, we did not find a significant association between colony area and remaining lifespan for *E. coli* (**Fig. 5f**), though we did find an association for *P. aeruginosa* infected animals (**Fig. 5g**). Both the number of animals with colonies and the variance of colony area within this group was very low across *E. coli* challenged animals in this set of experiments relative to *pmk-1* experiments despite using the same timing, which may provide a technical explanation for this disparity between experiments.

**Figure 5.**
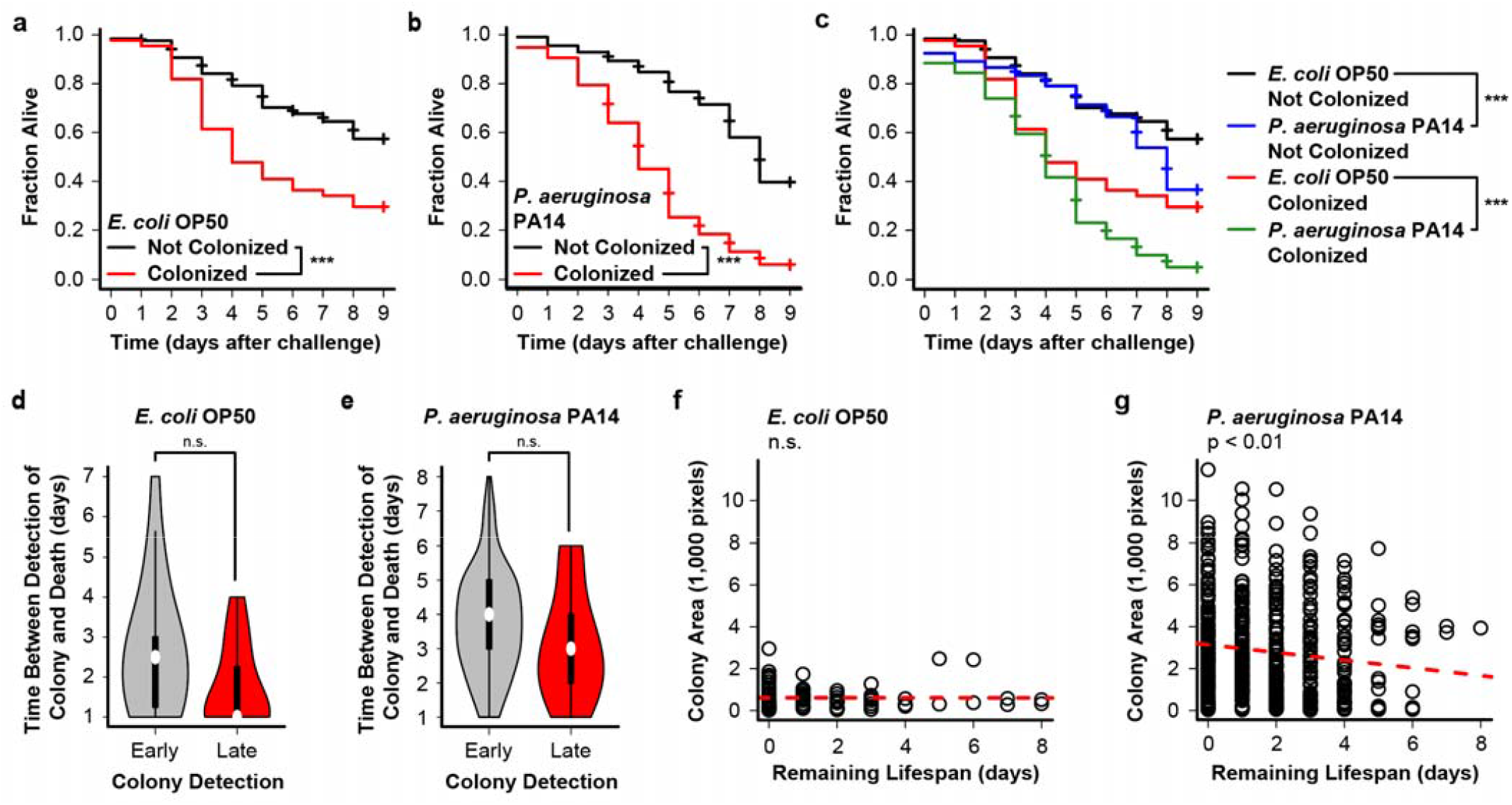
*C. elegans* challenged with *P. aeruginosa* PA14 are short-lived relative to animals challenged with *E. coli* OP50 in both colonized and uncolonized subgroups. *C. elegans* harboring colonies are substantially shorter lived than animals without colonies in both populations challenges with (**a**) GFP-labeled *E. coli* OP50 (p < 0.001, log rank test) and (**b**) mScarlet-labeled *P. aeruginosa* PA14 (p < 0.001, log rank test). (**c**) Lifespan of *P. aeruginosa* challenged *C. elegans* is significantly shorter than that of *E. coli* challenged animals in both colonized (p < 0.001, log rank test) and uncolonized (p < 0.001, log rank test) subpopulations. The time between initial detection of a colony and death of the animal trended lower but was not significantly different between animals with colonies that were detected late (day 3 or later post challenge) vs. early (day 1 or 2 post-challenge) for both animals challenged with (**d**) *E. coli* (p = 0.061, two-side Welch’s t test) and (**e**) *P. aeruginosa* (p = 0.11, two-side Welch’s t test). For *C. elegans* harboring an *E. coli* colony, the area of the colony on a given day (**f**) was not significantly associated with remaining lifespan for animals challenged with *E. coli* (p = 0.97, linear regression), but (**g**) was significantly and negatively associated with remaining lifespan for animals challenged with *P. aeruginosa* (p < 0.01, linear regression) animals. Each point represents one animal on one day. For violin plots, center bar or white point represent median, boxes represent upper and lower quartile, whiskers represent the 5^th^ and 95^th^ percentile, and points indicate outliers. Sample sizes: *E. coli* OP50, N_colonized_ = 40, N_uncolonized_ = 233, N_total_ = 273; *P. aeruginosa* PA14, N_colonized_ = 147, N_uncolonized_ = 127, N_total_ = 247. * p < 0.05, ** p < 0.01, *** p < 0.001, n.s. = not significant for indicated statistical test.

A further application of SICKO is to characterize colony dynamics in individual animals across the population. We asked whether progression of colony growth is linked to survival. We used the slope from a linear regression of colony area in individual animals over time as a first order estimate of the average colony growth rate for that animal. In both the wild type (**Fig. 6a**) and *pmk-1* (**Fig. 6b**) animals, we found that colony growth significantly and negatively correlated with survival among colonized individuals. Surprisingly, colony growth rate was similar between wild type and immune deficient *pmk-1* mutants (**Fig. 6c**). Colony growth rate also negatively correlated with survival for animals challenged with both *E. coli* (**Fig. 6d**) and *P. aeruginosa* (**Fig. 6e**). In the case of *P. aeruginosa*, we see that colony growth rate is higher, on average, than *E. coli* (**Fig. 6f**). This provides a more complete characterization of the colonization dynamics of *P. aeruginosa*. It not only colonizes animals at a higher rate than *E. coli*, but also grows faster once the colony is formed.

**Figure 6.**
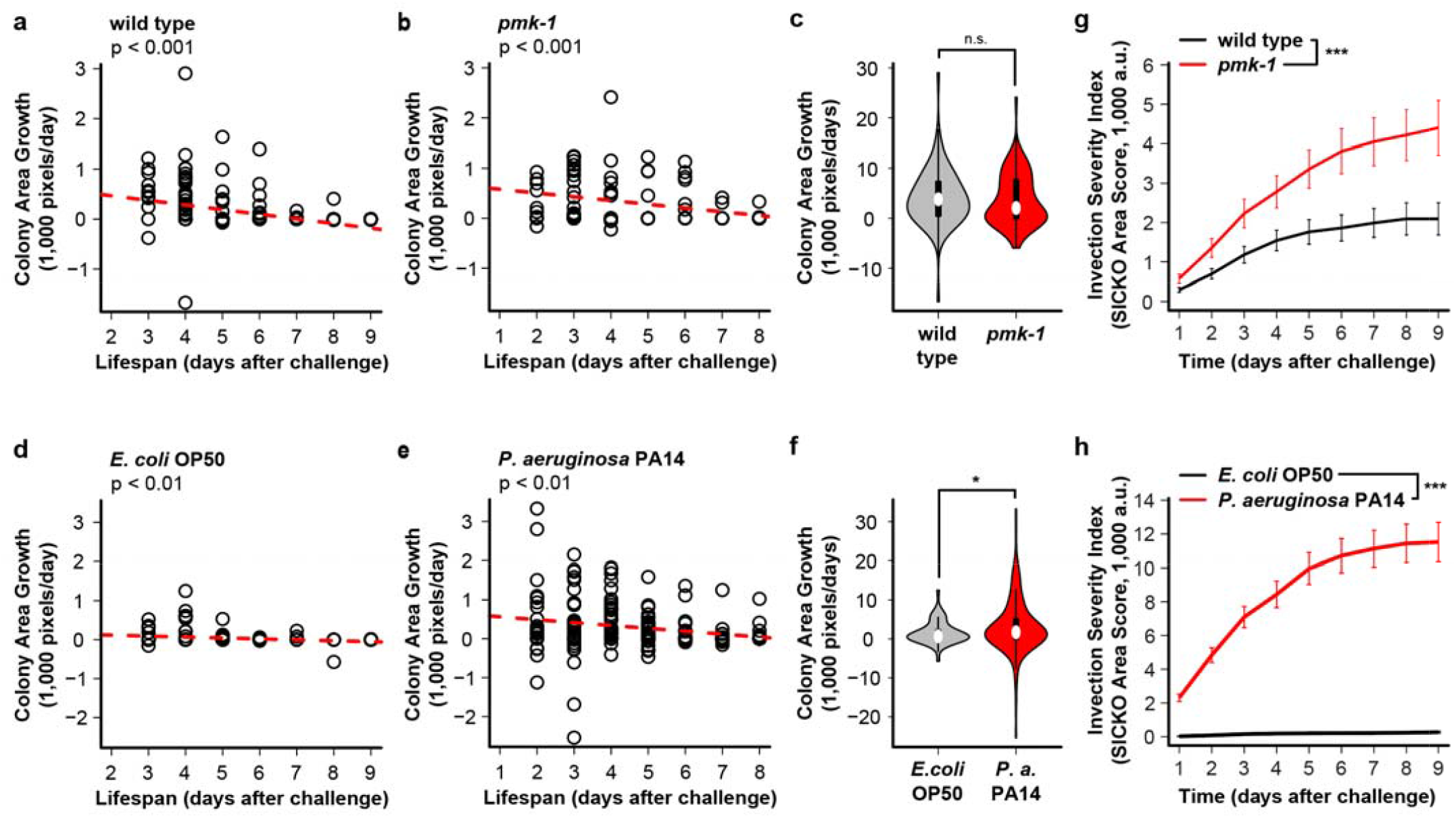
Faster colony growth is associated with reduced lifespan. Colony growth rate is significantly and negatively associated with lifespan in both (**a**) wild type (p < 0.001, linear regression) and (**b**) *pmk-1* (p < 0.001, log rank) animals. (**c**) Colony growth rate is not significantly different between wild type and *pmk-1* animals (p = 0.62, two-sided Welch’s t test). Colony growth rate is significantly and negatively associated with lifespan in *C. elegans* challenged with both (**a**) *E. coli* (p < 0.01, linear regression) and (**b**) *P. aeruginosa* (p < 0.01, log rank) animals. (**c**) Colony growth rate is significantly higher in *C. elegans* challenges with *P. aeruginosa* relative to *C. elegans* challenges with *E. coli* (p < 0.05, two-sided Welch’s t test). Individual colony growth rate was estimated as the slope of the colony area over time, calculated using linear regression; each point represents an individual animal (panels a-f). Infection severity is significantly higher for (**g**) *pmk-1* vs. wild type animals and for animals challenged with (**h**) *P. aeruginosa* vs. *E. coli*. Infection severity is estimated by adjusting the colony area in each animal for rate of colonization and prior deaths within the same treatment group using the SICKO coefficient (see **Methods**). For violin plots, center bar or white point represent median, boxes represent upper and lower quartile, whiskers represent the 5^th^ and 95^th^ percentile, and points indicate outliers. Sample sizes: wild type, N_colonized_ = 70, N_uncolonized_ = 171, N_total_ = 241; *pmk-1*, N_colonized_ = 69, N_uncolonized_ = 101, N_total_ = 170; *E. coli* OP50, N_colonized_ = 40, N_uncolonized_ = 233, N_total_ = 273; *P. aeruginosa* PA14, N_colonized_ = 147, N_uncolonized_ = 127, N_total_ = 247. * p < 0.05, ** p < 0.01, *** p < 0.001, n.s. = not significant for indicated statistical test.

### The SICKO score provides a metric of infection severity

The primary limitation of the SICKO system arises from the need to transfer animals away from the fluorescently labeled bacteria following exposure. This allows colonies established in the gut to be distinguished from the bacteria outside of the animal on the plate. During our validation studies, we found that the *pmk-1* and *P. aeruginosa* challenged animals were much more likely than wild type to die during the wash and transfer steps (**Fig. 1a[iii], Table 1**), introducing a high likelihood for selection bias. We speculate that the lost *pmk-*1 and *P. aeruginosa* animals were those with early severe colonies and that this may account, at least in part, for some of the unexpected differences observed in wild type vs. *pmk-1* animals, particularly in terms of the survival of animals with detectable infections (**Fig. 4c**). A related limitation is that no single metric provides a straightforward quantification of infection severity for pathogenic bacteria in a population at given point in time; in part because once an animal dies there is no longer a colony to quantify in terms of area or integrated intensity. To address these issues, we developed metric that reflects infection severity within a population, which we term the SICKO score, that modifies either the colony area or integrated intensity measurement for each animal with a weighting that reflects death prior to observation time (including animals lost during the wash and transfer steps), death during the observation time, and the proportion of the population afflicted by pathogen infection (see Methods for details). In essence, the SICKO score is an infection index that considers a bacterial colony of a given size more “severe” for bacteria if it causes more deaths and/or a higher colonization rate within the measured population. The *pmk-1* mutant animals showed a consistently higher SICKO score, calculated based on area of infection, throughout their lifespan relative to wild type animals (**Fig. 6g**). We observed a similar pattern when the SICKO score is derived from integrated infection intensity (**Fig. S2c**). The SICKO score was dramatically higher for animals challenged with *P. aeruginosa* vs. *E. coli*, reflecting the drastic difference pathogenicity between the two types of bacteria (**Fig. 6h**). In summary, the SICKO scores provide a composite metric of infection severity within a population for pathogenic bacteria.

**Table 1.** Animals lost during wash step. Counts of total live and dead *C. elegans* following the wash step (**Fig. a[iii]**) across each biological replicate experiment.

| Genotype | Bacteria | Replicate 1 |  | Replicate 2 |  | Replicate 3 |  | Combined |  |
| --- | --- | --- | --- | --- | --- | --- | --- | --- | --- |
|  |  | Alive | Dead | Alive | Dead | Alive | Dead | Alive | Dead |
| wild type | *E. coli* OP50 | 150 | 16 | 144 | 3 | 170 | 0 | 464 | 19 |
| *pmk-1* | *E. coli* OP50 | 160 | 50 | 95 | 10 | 132 | 0 | 387 | 60 |
| wild type | *E. coli* OP50 | 126 | 7 | 114 | 0 | 256 | 1 | 496 | 8 |
| wild type | *P. aeruginosa* PA14 | 130 | 50 | 229 | 2 | 191 | 10 | 550 | 62 |

## Discussion

Here we present SICKO, an imaging and analysis pipeline capable of monitoring fluorescently labeled bacteria colony progression over time in the tissues of individual free-crawling *C. elegans* on solid NGM media. Using the immune deficient *pmk-1* mutant, *E. coli*, and *P. aeruginosa* we validated the capacity of SICKO to quantify differences in dynamics of bacterial colonization and colony growth progression that would be arduous or impossible to reproduce using current standard cross-sectional approaches. We also developed the SICKO score, a composite metric of infection severity within a population of *C. elegans* challenged with a pathogenic bacterium that incorporates information on prior mortality and infectivity. Through longitudinal monitoring of bacterial colonization in individual animals, SICKO enables researchers to investigate in detail how the presence, severity, and dynamic changes in bacterial colonization impact survival.

Using SICKO, we found that colony progression does not appear to significantly different in *pmk-1* mutants challenged with *E. coli* when compared to wild type (**Fig. 2a**), but instead that *pmk-1* mutants had an increased susceptibility to colonization (**Fig. 2a, b, d**). This suggests that the shorter lifespan in *pmk-1* relative to wild type animals results from a higher fraction of the *pmk-1* animals harboring *E. coli* colonies, a subpopulation that is shorter lived in both *pmk-1* and wild type populations (**Fig. 4c**). Also surprisingly, the colonized *pmk-1* mutants appear to survive, if anything, slightly longer than colonized wild type animals (**Fig. 4c**), while uncolonized *pmk-1* animals had similar survival to uncolonized wild type animals, suggesting that *pmk-1* may not be a strong determinant of lifespan in the absence of pathogenic bacteria. We speculate that when comparing a wild type vs. *pmk-1* animal with a similar health status (aka similar expected remaining lifespan), the *pmk-1* animal is more likely to become colonized. Even with the capacity to separately examine survival in subpopulations with and without bacterial colonies, SICKO is unable to distinguish between (1) a model in which colonization causes reduced health and a shorter lifespan and (2) a model in which animals with a lower initial health status (and reduce expected lifespan) are more likely to become colonized. While these models are not mutually exclusive, because of this uncertainty in the direction of causality we urge caution in interpreting survival disparities in these datasets.

We also used SICKO to capture the higher colonization rate and negative health consequences for wild type *C. elegans* challenged with the strongly pathogenic *P. aeruginosa* PA14 relative to the weakly pathogenic *E. coli* OP50. Here we will highlight a technical challenge that arises when comparing distinct bacterial strains. We initially attempted to use the same GFP construct to label *P. aeruginosa* PA14 that we successfully used in *E. coli* OP50. However, we found that GFP expression was low in *P. aeruginosa* PA14, making colony detection difficult to quantify above background signal, particularly early or small colonies. We were able to successfully employ a similar expression construct using mScarlet in place of GFP. The background signal for mScarlet in *C. elegans* is much lower, allowing a comparable (though not identical) signal-to-noise ratio for *P. aeruginosa* PA14 mScarlet colonies relative to *E. coli* OP50 GFP colonies. Microscopy and image processing were calibrated to detect similarly sized colonies in worms challenged with both strains. When comparing bacteria strains, we also used colony area as a primary metric when comparing colonization between different bacteria to minimize artificial differences introduced by fluorophore expression level that can result from comparing integrated colony intensity. We note that differences in fluorophore expression between strains are difficult to eliminate and can influence some quantified characteristics of colony dynamics, such as the day when a colony becomes detectable. This is a limitation to keep in mind when comparing colonization parameters between different bacterial strains using SICKO or any other method that employs fluorophore-expressing bacteria to examine tissue colonization.

Finally, we noted the potential for survival bias that can arise later in life when there are disparities between comparison groups in the number of animals harboring colonies or the number of animals that died prior to an observation time. For example, a highly pathogenic strain of bacteria that causes animal death earlier or at a lower bacterial load than a less pathogenic bacteria may artificially appear to cause a less severe infection of colony area alone is used as a metric, simply because animals with the most severe infection died at an earlier timepoint. To account for this complication, we developed the SICKO score as a composite metric of infection severity that modifies colony area or intensity based on earlier death and colonization rates in the population. We find that this allows both *pmk-1* to be better distinguished from wild type animals (**Fig. 6g**) and *P. aeruginosa* challenged animals to be better distinguished from *E. coli* challenged animals (**Fig. 6h**). We note that this score increases with higher colonization potential, and is therefore only conceptually applicable to pathogenic microbes (in which colonization is assumed to negatively impact host health). While our original motivation was to study pathogenic bacteria, SICKO is also compatible with studying beneficial bacteria. In this case all metrics, with the exception of the SICKO score, remain valid. An alternative index for beneficial bacteria will need to be developed and validated for this application.

Because many components of the SICKO system analysis have been automated, any researcher with moderate *C. elegans* handling experience and minimal specialized equipment should be capable of utilizing the system and producing rich characterization of bacterial colonization phenotypes in *C. elegans* in a wide range of test conditions. SICKO is compatible with mutant and transgenic strains, RNAi, single microbes, combinations of microbes labeled with different fluorophores, and variable environmental conditions. When combined with emerging automated image collection systems, SICKO will open the door to high-throughput analysis of host-microbe interaction. As noted above, SICKO can be used to examine colonization dynamics for beneficial bacteria and how these dynamics impact different aspects of *C. elegans* health. Currently, one major limitation of SICKO lies in the washing step and the use of antibiotics. In the present study, the animals are initially grown on live unlabeled *E. coli* OP50 prior to challenge with fluorescently labeled *E. coli* or *P. aeruginosa*, leaving open the possibility that some animals are acquiring unlabeled *E. coli* colonies prior to challenge that are influencing health but are not being detected. Gentamycin is included in the single-worm environments to both prevent growth of the unlabeled *E. coli* food source and select for the plasmid in fluorescently labeled bacteria colonizing the animal. In early testing of SICKO, we tested animals exposed to fluorescently labeled *E. coli* OP50 and observed very low rates of colonization, so we speculate that this is not a major issue. However, in the future we may improve this step by using non-living food sources, such as heat, UV, or PFA killed bacteria, or axenic food both before and after challenge with the fluorescent microbe of interest. It may also be feasible to create a biosensor system in which only colonizing bacteria produce fluorescent (perhaps through detection of contact with the *C. elegans* intestine or intestinal pH). This would eliminate the need for a wash step and allow worms to be monitored immediately following microbial exposure. This solution is of interest but will require substantial technical development and validation. To conclude, the SICKO system allows an avenue for detailed testing of novel therapies to combat pathogens, conduct mechanistic studies of innate immune function, and explore the influence of both pathogenic and beneficial bacteria on host health and survival.

## Methods

### *C. elegans* strains and maintenance

*C. elegans s*train KU25 (*pmk-1(km25) IV*) was obtained from the *Caenorhabditis* Genetics Center (CGC), which is funded by NIH Office of Research Infrastructure Programs (P40 OD010440). Wild type (N2) worms were originally obtained from Dr. Matt Kaeberlein (University of Washington, Seattle, WA, USA). *C. elegans* were cultivated on 60 mm culture plates containing nematode growth media (NGM) solidified with agar as previously described (Sutphin and Kaeberlein, 2009). Briefly, NGM agar plates were spotted with 300 µl of the *E. coli* strain OP50 grown overnight in lysogeny broth (LB) in a 37 °C orbital shaker and allowed to dry for a minimum of 16 hrs. Worms were maintained with abundant food for a minimum of three generations following recovery from frozen stock or from dauer larvae prior to running experiments. Animals were incubated at 20 °C and passed every 3-4 days to fresh NGM plates containing food.

### Bacteria strain construction

*E. coli* containing the pMF230 plasmid, which includes both an ampicillin resistance cassette and eGFP driven by a constitutively active trc promoter, was obtained from Addgene (catalog #62546) and cultivated on LB agar plates containing 100 μM ampicillin to select the plasmid. A gentamycin resistance cassette from plasmid pFGM1 (Addgene catalog #64949) was and cloned into pMF230 plasmid, creating pMFG-eGFP, a constitutively active eGFP plasmid with 10 mg/ml gentamycin resistance. Plasmids were isolated with a QIAprep Miniprep Kit (Qiagen) and quantified using 260 nm/280 nm absorbance on a Biotek Synergy H1 plate reader. OP50 was made competent and transfected with pMFG-eGFP resistance plasmid as previously described (Choi et al., 2006). Bacteria colonies displaying green fluorescence were selected by plating LB supplemented with 10 mg/ml gentamycin. We constructed plasmid pMFG-mScarlet by cloning mScarlet into plasmid pMFG-eGFP to replace eGFP. *P. aeruginosa* strain PA14 was made competent and transfected with plasmid pMFG-mScarlet as previously described (Choi et al., 2006).

### Validation experimental design and replication

In this study we report two sets of experiments. The first set compares wild type (strain N2) worms vs. *pmk-1(km25)* loss of function mutants (strain KU245) challenged with GFP-labeled *E. coli* OP50. The second set compares wild type worms challenged with GFP-labeled *E. coli* OP50 to mScarlet-labeled *P. aeruginosa* PA14. Each set consisted of three independent biological replicates. All figures present the pooled data across all three replicates within the indicated experiment set. The number of animals lost during the wash step are provided in **Table 1**, and the number of animals examined during the observation step are provided in **Table 2** for each replicate and for the pooled dataset.

**Table 2.** Sample size per replicate. Counts of total, colonized, and uncolonized animals in each experimental replicate.

|  |  | Replicate 1 |  |  | Replicate 2 |  |  | Replicate 3 |  |  | Combined |  |  |
| --- | --- | --- | --- | --- | --- | --- | --- | --- | --- | --- | --- | --- | --- |
| Genotype | Bacteria | Colony | No Colony | Total | Colony | No Colony | Total | Colony | No Colony | Total | Colony | No Colony | Total |
| wild type | *E. coli* OP50 | 53 | 30 | 83 | 5 | 65 | 70 | 12 | 76 | 88 | 70 | 171 | 241 |
| *pmk-1* | *E. coli* OP50 | 23 | 17 | 40 | 21 | 51 | 72 | 25 | 33 | 58 | 69 | 101 | 170 |
| wild type | *E. coli* OP50 | 12 | 74 | 86 | 19 | 72 | 91 | 9 | 87 | 96 | 40 | 233 | 273 |
| wild type | *P. aeruginosa* PA14 | 50 | 41 | 91 | 72 | 17 | 89 | 52 | 42 | 94 | 147 | 127 | 274 |

### Preparation of single-worm culture environments

Single worm culture environments were prepared as previously described (Espejo et al., 2022). Briefly, for each environment a Terasaki tray is mounted inside a single-well tray using a custom printed 3D adapter. The plastic interior surface of the Terasaki tray surrounding the wells is coated with an aversive barrier by applying a solution of 10 mg/mL palmitic acid (to prevent fleeing) 40 units/mL nystatin (to prevent fungal contamination) in 30% Tween-20 35% ethanol and allowing the liquid to fully evaporate. Each well of the Terasaki tray is filled with NGM solidified with low-melt agarose in place of agar (lmNGM), supplemented with 1 mg/mL gentamycin to select fluorescent plasmids, and seeded with *E. coli* OP50 grown in LB overnight in a 37 °C shaker, pelleted, and resuspended at a 10-fold concentration in 85 mM NaCl solution. The space surrounding the Terasaki tray in the single-well tray is filled with saturated water-absorbing crystals to provide humidity. The single-well tray is closed and sealed with parafilm until worms are ready to load (no more than 24 hours).

### Bacterial challenge, wash, and plating

*C. elegans* were age synchronized by hypochlorite treatment (Porta-de-la-Riva et al., 2012). and plated on to NGM seeded with *E. coli* strain OP50. For worms challenged with *E. coli*, L4 larval stage worms were transferred to NGM plates supplemented with 500 μM floxuridine (FUdR) to prevent reproduction and 1 mg/ml gentamycin for plasmid selection and spotted with GFP labeled *E. coli* OP50 bacteria and incubated at 20 °C for 7 days. For worms challenged with *P. aeruginosa*, L4 larval stage worms were first transferred to NGM plates supplemented with 500 μM floxuridine (FUdR) to prevent reproduction seeded with unlabeled *E. coli* OP50 and incubated for 5 days. These day 5 adult animals were then transferred to NGM plates supplemented with 500 μM floxuridine (FUdR) to prevent reproduction and 1 mg/ml gentamycin for plasmid selection and spotted with mScarlet labeled *P. aeruginosa* PA14 bacteria and incubated at 20 °C for 2 days.

In both cases, worms were transferred following challenge (day 9 of adulthood) were transferred to fresh NGM plates supplemented with 500 μM FUdR and seeded with unlabeled *E. coli* OP50. Worms were placed on the edge of the plate outside of the bacterial spot prompting them to crawl toward the food. The animals are incubated for at least 16 hours at 20 °C to allow non-adherent GFP-expressing bacteria to fully pass out of their gut and be shed from the external surface of their body. Following this process, the only remaining GFP-expressing bacteria are in adherent colonies in the *C. elegans* gut. At the end of the 16-hour incubation, randomly selected animals are transferred to individual wells of prepared single-worm culture environments. The number of animals transferred, the number remaining alive on intermediate plate, and the number that died on the plate are all recorded for calculation of SICKO coefficient later.

In optimizing SICKO, we examined multiple exposure windows for wild type *C. elegans* to GFP-expressing *E. coli* OP50 and mScarlet-expressing *P. aeruginosa* to optimize the challenge step. Wild type animals challenged for with *E. coli* OP50 for 7 days starting at the L4 larval stage, or with *P. aeruginosa* for 2 days starting at day 5 of adulthood, result in detectable gut infections in approximately 30-50% of animals over the subsequent observation period in test experiments. Exposure to either strain during development or for fewer days starting from the L4 stage results in colonies only rarely (data not shown).

### Image collection and processing

Starting on the day the animals are loaded onto the single-worm culture environments, three images of each animal are captured daily using the GFP channel (excitation: 425/60 nm, emission: 480 nm) of a Leica M205 FCA Fluorescent Stereo Microscope equipped with a Leica K6 sCMOS monochrome camera using 2.5x zoom and saved as TIF files in the folder structure specified by the SICKO software documentation (**Fig. S3**) (Freitas, 2024). Capturing multiple images per animal ensures the signal is not impacted by worm movement (i.e., a non-blurry image can be selected for each animal). Worms are exposed to blue light to stimulate movement and recorded as dead if no movement observed, or fled if the animal is no longer present in the well.

### Image analysis and output

Version March.13.2024.v1 of the SICKO software, built in MATLAB version 2022b, was used in this study (Freitas, 2024). SICKO takes as input raw fluorescent images each containing a single worm infected with fluorescently labeled bacteria, removes background, identifies the infection area, and quantifies the area and integrated fluorescent intensity of the infection. The SICKO image processing GUI allows users to manually select image regions that contain artifacts (usually fluorescently labeled bacteria outside of the worm or light pollution) that are removed from analysis. The SICKO analysis software can be downloaded along with complete documentation at: https://github.com/Sam-Freitas/SICKO (Freitas, 2024). A step-by-step protocol for using the software is provided. Generation of figures and statistics using the SICKO output files was conducted in RStudio version 2023.12.1 Build 402 running R version 4.3.2. The scripts used to generate these figures are available with instruction for use at: https://github.com/lespejo1990/SICKO_Analysis (Espejo, 2023).

SICKO analysis relies on a threshold to distinguish infection from background. A user defined threshold determines a mask that captures the area in the image representing bacterial colony (the “colony mask”). To compensate for background differences across the field of view, background correction is performed radially of the colony mask, ensuring animal position within the field of view minimally impacts signal intensity and accounts for uneven background. The intensity of all pixels within the colony mask is integrated to calculate the integrated infection intensity for each animal. The number of pixels within the colony mask defines the colony area.

Examining the area or integrated intensity of a colony over time in live animals across a population provides information on the state of the colony size at a given time point but has limited utility in tracking colony progress over time because it cannot account for animals that died during the pathogen challenge, during the wash, or during an earlier observation time. Animals that died at earlier stages likely represent a more severe response to colonization, and thus simply quantifying infection area or integrated intensity at a given time point will tend to underestimate the pathogen severity by excluding data from dead animals. We developed an infection severity index for pathogenic bacteria called the SICKO score that provides a quantitative metric of pathogen colony (aka “infection”) progress within a population that systematically accounts for the number of worms that died before a given time, including during the post-challenge wash, worms that died at the time of observation, and fraction of animals harboring a colony within a population (a metric of infectivity). To determine the SICKO score, we first calculate a SICKO coefficient, which is then used to weight the colony area or integrated intensity for each animal at each time point. The SICKO coefficient is updated for each time point to reflect the fraction of animals that died with a colony up until that day.

First, on each day of observation, *t*, we calculate the cumulative number of animals that have died while harboring a bacterial colony during the observation period (**Fig. 1a[v]**) up until that day, *D*_*CO*_*(t)*, and the total number of animals that died while harboring a bacterial colony across the full observation period, 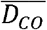:

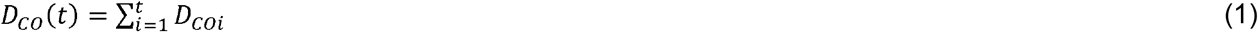

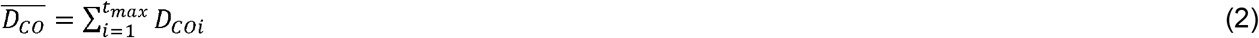

where *D*_*COi*_ is the number of worms the died while harboring a colony at time point *i*, and *t*_*max*_ is the last day of observation.

Next, we estimate the number of worms that died during the challenge **Fig. 1a[ii]** and wash **Fig. 1a[iii]** steps while harboring a bacterial colonization, *D*_*CW*_:

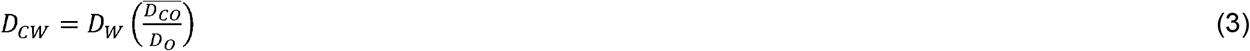

where *D*_*W*_ is the total number of worms that died during the challenge and wash steps, *D*_*CO*_ is the number of animals that died while harboring a colony during the full observation period (**Fig. 1a[iv]**), and *D*_*O*_ is the total number of worms that died during the observations period (**Fig. 1a[iv]**). We note that this cannot be measured directly, because while worms are on the challenge and wash plates, they will have fluorescent bacteria in their intestine and on their external surface that is not part of an established tissue colony but cannot be distinguished from bacteria in colonies. In our case, we make the assumption that the fraction of worms that die with a colony during the challenge and wash steps will be the same as the fraction that die with a colony during observation period. Other assumptions and approaches can be employed in this case, for example that all worms lost during challenge and wash harbor a colony (which may be warranted for highly pathogenic bacteria that are likely to result in early animal death).

From equation (3), we next estimated number of worms that ever had a detectable bacteria colony at any time during the experiment, *P*_*C*_:

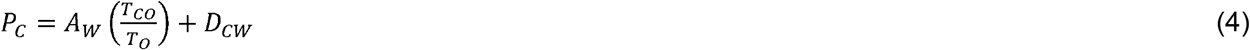

where *A*_*W*_ is the number of animals alive after the challenge and wash step (including those not selected for observation, **Fig. 1a[iii]**), *T*_*CO*_ is the total number of non-censored animals that were every observed to have a colony in the observation group (**Fig. 1a[v]**), and *T*_*O*_ is the total number of non-censored animals in the observation group (**Fig. 1a[v]**).

Using the values from Equations (1), (3), and (4), we next calculate the SICKO coefficient, *S*_*C*_ which provides a population-level weighting that increases with increasing infectivity (fraction of animals the develop a colony) and toxicity (fraction of animals that die with a colony prior up until the current day, *t*):

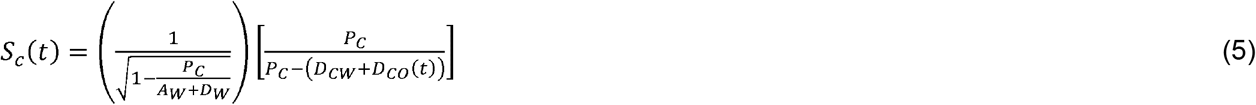

In Equation (5), the term in the left parentheses represents a weighting that increases with the fraction of animals that develop a bacterial colony, while the term in the right brackets represents a weighting that increases with the number of animals that have died while harboring a colony up until the current day, *t*.

Finally, the SICKO coefficient in Equation 5 is used to as a weight to calculate at SICKO infection severity index based on either the measured colony area, *S*_*I*,*area*_*(t)*, or integrated intensity, *S*_*I*,*intensity*_*(t)*, for each worm, *w*, on each day, *t*:

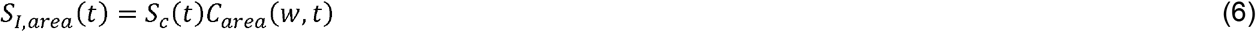

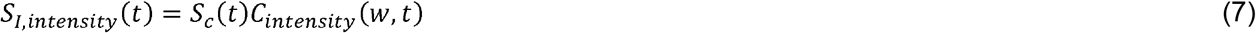

where *C*_*area*_*(w*,*t)* and *C*_*intensity*_*(w*,*t)* are the measured colony area and integrated intensity, respectively, for worm *w* on day *t*. The values for *S*_*I*,*area*_ and *S*_*I*,*intensity*_ are plotted in **Fig. 6e,f** and **Fig. S2c**, respectively.

## Data availability

All data needed to evaluate the conclusions in the paper are presented in the paper and Supplementary Materials. The datasets generated and used during the current study are publicly available through the ReDATA repository at the University of Arizona (DOI: 10.25422/azu.data.25749384, DOI: 10.25422/azu.data.25749372). The SICKO image processing (version March.13.2024.v1) (Freitas, 2024) and analysis (version December.15.2022.v1) (Espejo, 2023) code is publicly available for download.

## Acknowledgements

This research was funded by NIH/NIGMS grant R35GM133588 to G.L.S.

## Author Contributions

L.S.E., S.F., and G.L.S. initially conceived of and developed the conceptual ideas behind SICKO. S.F. and V.H. developed the image processing software. L.S.E., S.F., and V.H. developed the analysis software. L.S.E. and H.D. created novel plasmids and bacteria strains. L.S.E., L.C., A.A., J.B., A.H., D.D., S.H., and D.K. conducted primary experimental planning and data collection. L.S.E., V.H., L.C., and A.A. conducted data analysis. H.D. provided training and experimental support. G.L.S. supervised all aspects of this work. L.S.E., S.F., V.H., and G.L.S. drafted the manuscript. All authors edited and approved the manuscript.

