## Supplemental Figures for "SICKO: Systematic Imaging of *Caenorhabditis* Killing Organisms"

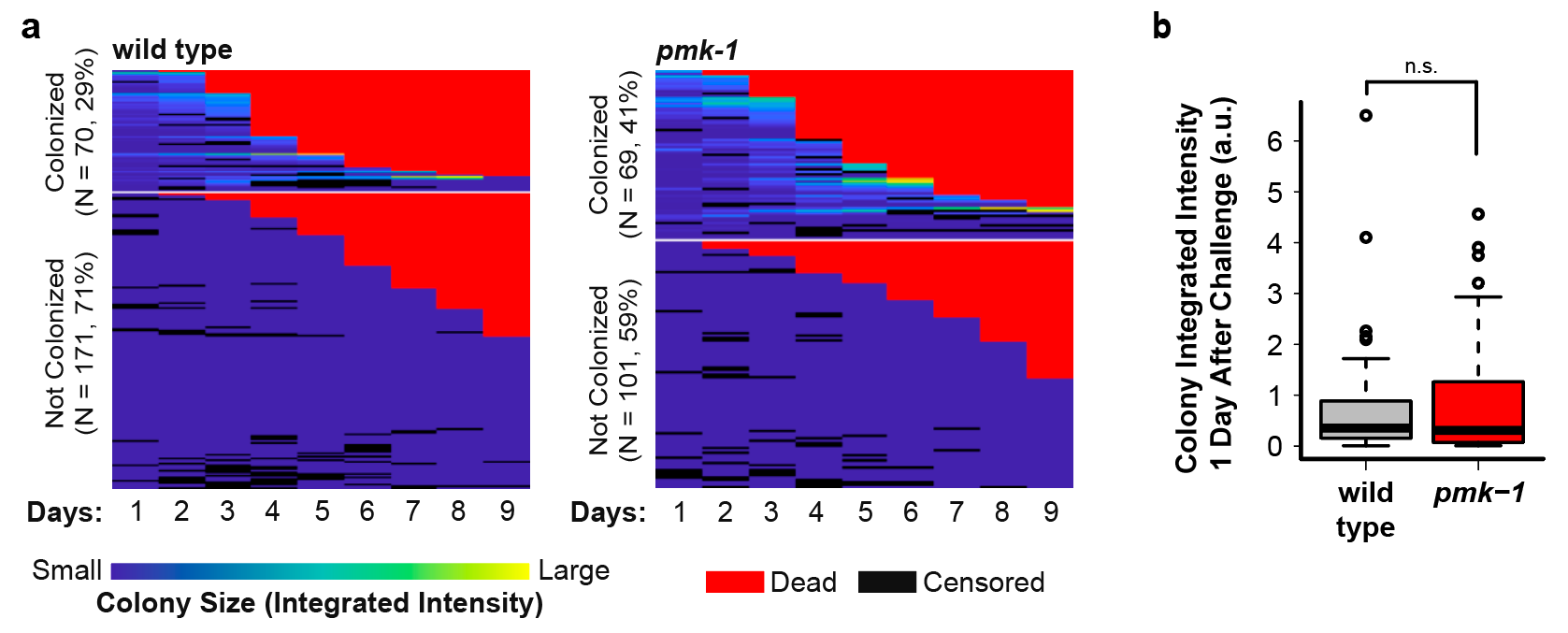

**Figure S1. *C. elegans* lacking *pmk-1* are more susceptible to *E. coli* colonization than wild type.** (**a**) Heatmap representing colony size (based on integrated fluorescence intensity) for wild type (left) and *pmk-1* knockout (right) *C. elegans* challenged with GFP-labeled *E. coli*. Death was highly correlated with colonization status in both wild type (p < 0.001, Pearson’s chi-squared test) and *pmk-1* (p < 0.001, Pearson’s chi-squared test) animals, and with genotype (p < 0.01, Pearson’s chi-squared test). (**b**) The mean integrated fluorescence intensity (p = 0.53, two-sided Welch’s t test) is not significantly different between wild type and *pmk-1* knockout *C. elegans* 1 day following challenge. For box-and-whisker plots, center bar or white point represent median, boxes represent upper and lower quartile, whiskers represent the 5^th^ and 95^th^ percentile, and points indicate outliers. Sample sizes: wild type, N_colonized_ = 70, N_uncolonized_ = 171, N_total_ = 241; *pmk-1*, N_colonized_ = 69, N_uncolonized_ = 101, N_total_ = 170. * p < 0.05, ** p < 0.01, *** p < 0.001, n.s. = not significant for indicated statistical test.

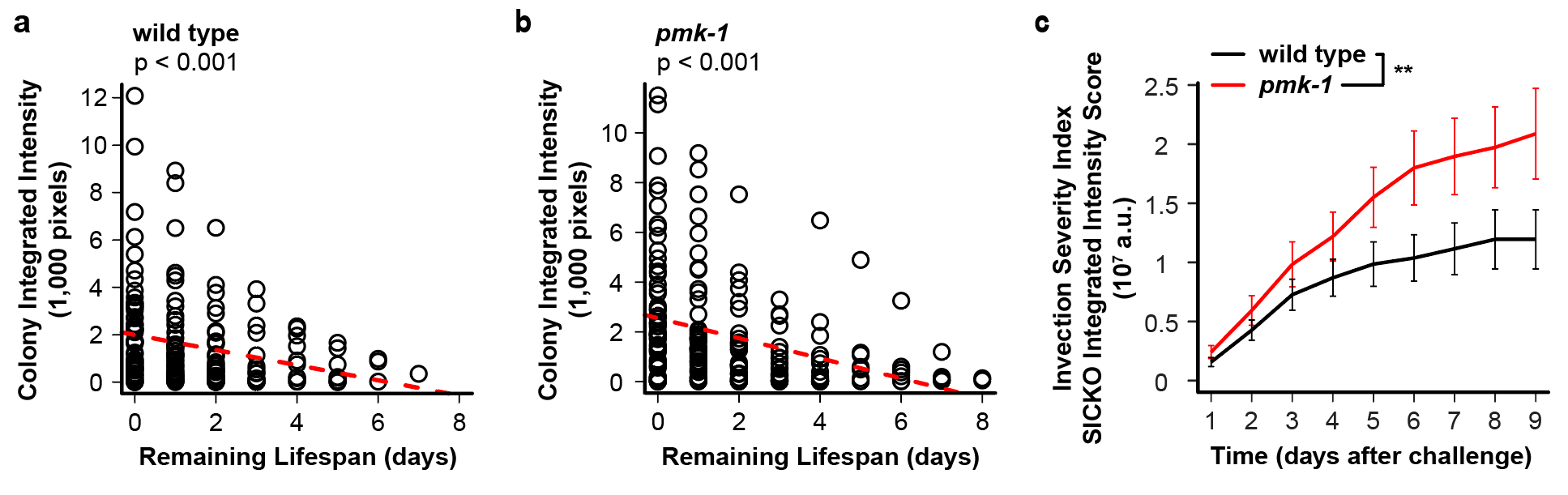

**Figure S2. Colony integrated fluorescence intensity is associated with remaining lifespan.** For *C. elegans* harboring an *E. coli* colony, the integrated fluorescence intensity of the colony on a given day was significantly and negatively associated with remaining lifespan in both (**a**) wild type (p < 0.01, linear regression) and (**b**) *pmk-1* (p < 0.001, linear regression) animals. Each point represents one animal on one day. (**c**) Infection severity is significantly higher for *pmk-1* vs. wild type animals. Infection severity is estimated by adjusting the colony integrated fluorescence intensity in each animal for rate of colonization and prior deaths within the same treatment group using the SICKO coefficient (see **Methods**). Sample sizes: wild type, N_colonized_ = 70, N_uncolonized_ = 171, N_total_ = 241; *pmk-1*, N_colonized_ = 69, N_uncolonized_ = 101, N_total_ = 170. * p < 0.05, ** p < 0.01, *** p < 0.001, n.s. = not significant for indicated statistical test.

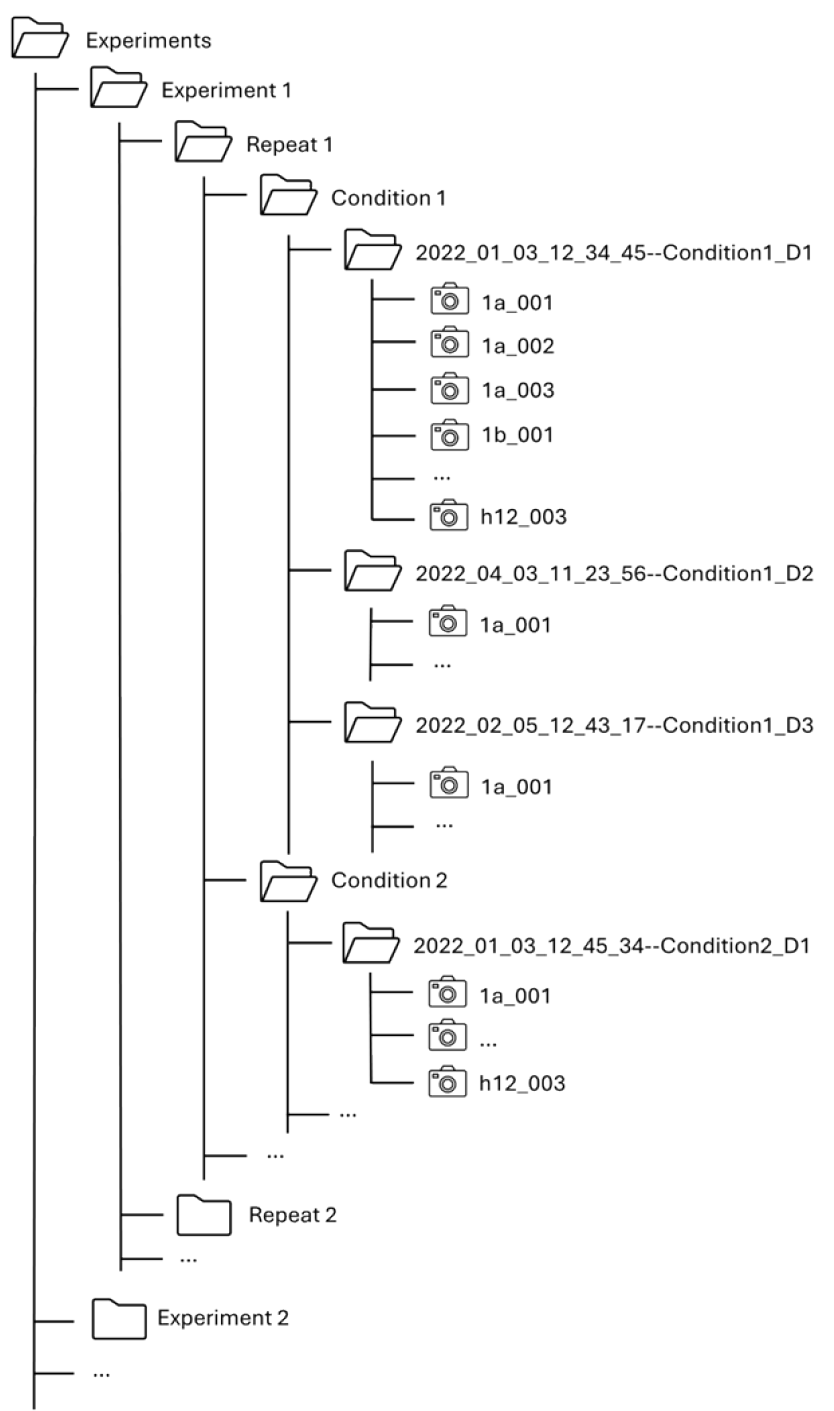

**Figure S3. Required folder structure for SICKO image processing.**
